# Membrane Remodeling by the Collective Action of Caveolin-1

**DOI:** 10.1101/2025.10.03.680304

**Authors:** Korbinian Liebl, Gregory A. Voth

## Abstract

Caveolin-1 proteins scaffold 50-100nm large invaginations in the plasma membrane to mediate critical cellular processes. As revealed recently by cryo-electron microscopy, several caveolin-1 protomers can fold into a disk-like structure that embeds in the cytoplasmic leaflet. This 8S complex represents a basal component to drive membrane curvature via higher-order interactions. The biophysical mechanisms behind the membrane remodeling, however, have remained elusive. To address this shortcoming, we have developed a new bottom-up coarse-grained model to overcome the substantial computational limitations for this large system. During simulations with the coarse-grained model, the complexes increasingly coordinate as partially mediated by attractive electrostatic interactions between scaffolding domains. The coordination of complexes strongly correlates with membrane protrusion, as approaching complexes amplify localized stress in the exoplasmic leaflet. Thus, proximity of two CAV1-8S complexes induces dynamic curvature generation that can facilitate access for signaling partners. This mechanism is further explored in clusters of multiple CAV1-8S complexes that form large-scale membrane invaginations.

## Introduction

Many cellular functions require remodeling of the membrane shape or composition. This behavior is strikingly manifested in the biogenesis of caveolae where caveolin proteins together with other cofactors (e.g., cavin) generate 50-100nm large plasma membrane invaginations and mediate various cellular processes including mechanoprotection, lipid homeostasis, and endocytosis.^1–8^ Dysregulation of caveolin-1 is hence implicated in several diseases such as cardiovascular hypertrophy, cancer, lipodystrophy and diabetes.^5, 9–13^

Caveolae assembly and function have been characterized with high-resolution imaging techniques as well as lipid-binding and membrane tension assays, showing that caveolin-1 (CAV1) proteins cluster cholesterol at the plasma membrane and drive membrane curvature even in the absence of cavin.^14–21^ Binding by cavin, however, further stabilizes and compacts the initial caveolae architecture.^20, 22, 23^ These assemblies fulfill versatile and contrasting roles, regulated by the concentration of EHD2 and other recruited components: Caveolae assemblies act as buffers to membrane tension but can also bud off from the plasma membrane, a critical step in endocytosis.^8, 17, 20, 22, 24^

Despite the extensive characterization of CAV1 function, the biophysical basis underlying these processes has largely remained elusive. In particular, the initial flat-to-curved transition of the plasma membrane induced by CAV1 is still an enigma^25^ that we address in this study. The understanding of caveolae is thus still in its infancy, mainly because its structural basis has been determined only recently. Using cryo-electron microscopy (cryo-EM), it has been resolved that eleven CAV1 protomers assemble into a disc-like structure (~14nm diameter) with a central beta-barrel formed by the C-termini of the protomers.^26–28^ The hydrophobic profile of this CAV1-8S complex suggests a monotopic arrangement (embedding only in the cytoplasmic leaflet) in which the beta-barrel remains solvent-exposed (Fig. 1A). This leaflet replacement model has been further supported by Langmuir film-balance measurements.^29^ To get a better understanding of the lipid-CAV1-8S interactions, various groups have studied this system with both all-atom (AA) and coarse-grained (CG) molecular dynamics (MD) simulations.^29–32^ All-atom MD simulations indicate that the CAV1-8S complex extracts cholesterol from the lipid membrane and accommodates it in its beta-barrel, whereas cholesterol accumulation at the distal leaflet is likely enhanced by post-translational modifications.^30^ Moreover, these AA MD simulations indicate that a single CAV1-8S complex induces only slight, local bending in the exoplasmic leaflet, hence suggesting a collective action that involves several complexes to remodel the membrane shape.^29, 30^

**Fig. 1.**
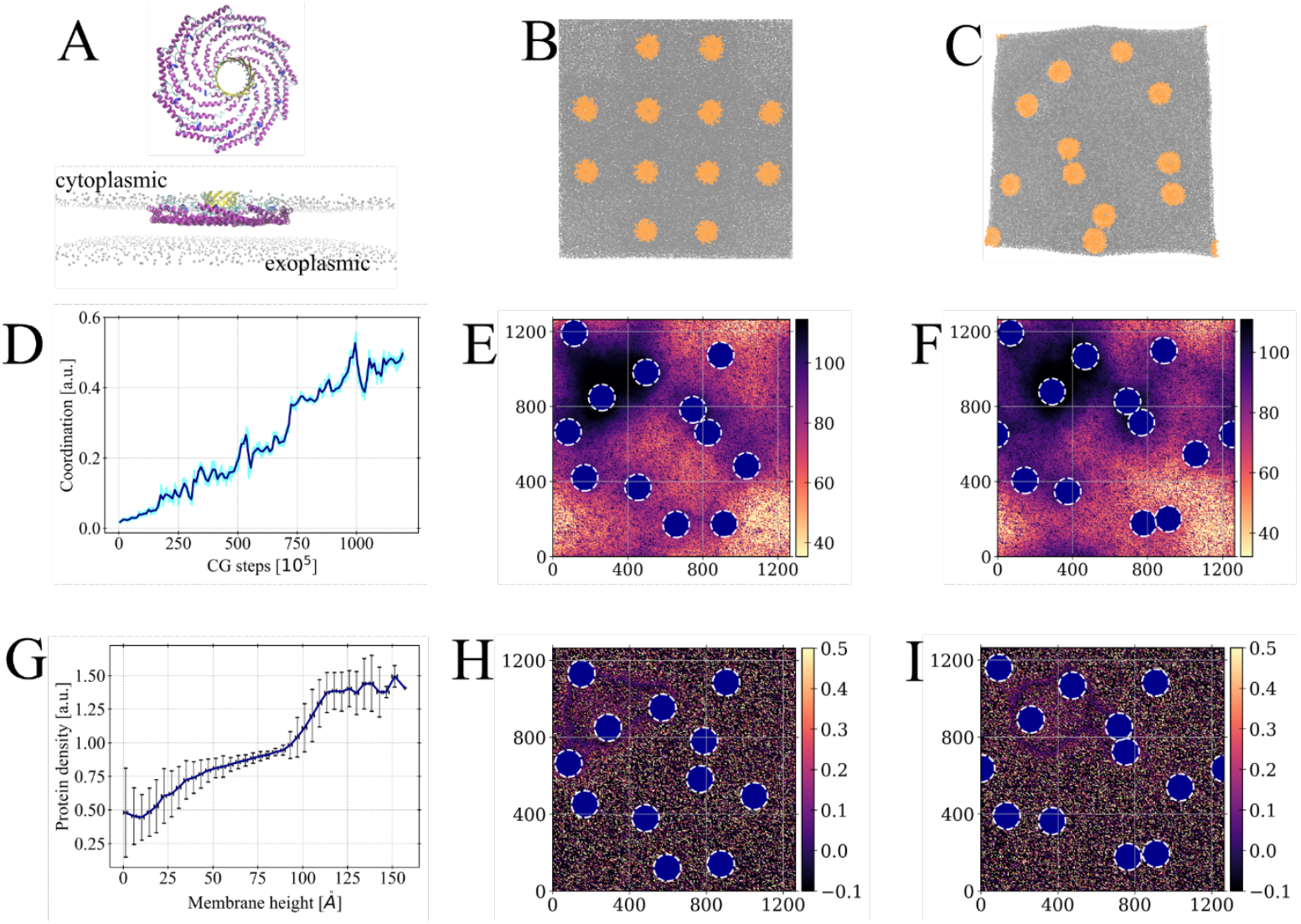
Coordination and collective membrane remodeling by the CAV1-8S complexes. **(A)** Structure of the CAV1-8S complex shown in top and side view. Note that the complex is embedded in the cytoplasmic leaflet. **(B)** Starting configuration for a large-scale CG simulation that includes 12 complexes (shown in orange). **(C)** Final structure of the simulation that indicates substantial diffusion and pairing of the CAV1-8S complexes. **(D)** Coordination of the complexes increases steadily during simulation which quantifies the tendency to cluster formation. **(E, F)** Height-profiles (units of Å) of the cytoplasmic leaflet computed via Gaussian Process Regression for representative frames of the simulation. Blue circles with dashed-white contour mark the positions of the complexes. **(G)** Positive correlation between membrane height and protein density quantifies curvature sensing and generation by the CAV1-8S complexes. Larger membrane protrusions arise only in the presence of multiple complexes. **(H, I)** Mean curvature profiles (units of Å^−1^) for representative frames derived from Gaussian Process Regression. Purple contours around (400,1000) highlight positive mean curvature and hence formation of a larger invagination of the membrane.

Accurate simulations of such large-scale processes reach the limits of current state-of-the-art computational hardware. However, by developing a bottom-up coarse-grained (CG) model for the CAV1-8S complex embedded in a heterogeneous lipid bilayer, we can overcome these substantial limitations. Importantly, as explained later, our CG model is also transferable to (self-assembling) lipid-bilayer systems without CAV1 and is solvent-free. Thus, it is scalable beyond 100nm and allows us to track diffusion of the complexes on similar length-scales.

Simulations with this CG model permit us to monitor coordination of the complexes. By leveraging Bayesian machine learning techniques, we also show that membrane protrusion and curvature arise due to the collective action of different CAV1-8S complexes, not from a single one. The approaching complexes sense each other’s local stress in the exoplasmic leaflet, causing pronounced tilting of the CAV1-8S disks in the near-distance regime. Interestingly, such a tilting or dynamic curvature mechanism has recently been hypothesized to regulate signaling pathways such as eNOS.^33^ Furthermore, we suggest that caveolae assembly may be promoted by the distinctive electrostatic profiles of the CAV1 scaffolding domains, characterized by four particular amino acids. The sequence of CAV1 thus not only encodes self-assembly into the 8S-oligomers, but also specific higher-order interactions that drive clustering of the CAV1-8S complexes.

In addition, the CG-model enables us to sample formation of ~70nm broad invaginations in the membrane at CAV1-cluster sites and hence to obtain molecular insight into this initial scaffolding-step of caveolae biogenesis. Recorded energy profiles show a minimum at intermediate invaginations, and further protrusion leads to a reduction in the high-l modes, indicating a loss of polyhedral symmetry. Our simulations hence suggest that tighter, more restricted invaginations need additional cofactors to sustain an ordered symmetric pattern of the CAV1-8S induced invaginations.

Thus far, accurate large-scale simulations of the embedded CAV1-8S complex have turned out to be a major challenge, and top-down CG models such as Martini can induce nonphysical membrane curvature for this system.^30^ In this work, we have therefore employed a new approach that allows for accurate and scalable CG simulations of this system and that is likely transferable to a broader class of protein-membrane systems. Application of our model offers new insight into fundamental biophysical behavior of CAV1-8S complexes which is of high relevance for our understanding of caveolae assembly and associated signaling pathways.

## Results

### Dynamic membrane remodeling and coordination of CAV1-8S complexes

Utilizing bottom-up techniques, we have trained a coarse-grained model that captures the monotopic and hence highly asymmetric embedding of the CAV1-8S complex in the heterogeneous lipid bilayer (70/30% POPC/CHOL, see Supporting Information Movie S1). Note that our CG model is significantly coarser than commonly used top-down CG-models such as Martini, especially since it is solvent-free (Fig. S1 of the SI). We emphasize that solvation effects are not neglected but intrinsically incorporated in our CG interactions that represent potentials of mean force between CG sites. Moreover, the lipid parameterization accurately describes self-assembly, bending modulus, and distribution functions of symmetric lipid bilayers (Movie S2, Figs. S2-S4). For these reasons, our CG-model is transferable and highly scalable, enabling the systematic simulation of membrane systems larger than 100nm that include multiple CAV1-8S complexes. Thus, we harness the power of this model to study membrane remodeling on scales larger than even the typical size of caveolae assemblies.

As a first step, we have simulated 12 CAV1-8S complexes embedded in a large ~125×125nm^2^ membrane bilayer (Fig. 1). During the simulation, the protein complexes show substantial diffusion along the cytoplasmic leaflet and increasingly interact with each other (Figs 1B-D, S5). Intriguingly, distinct complexes can approach each other to form clusters, and the membrane is partitioned into CAV1-rich and CAV1-depleted regions. Based on the recorded statistics we investigated the relationship between CAV1-density and membrane topology. To obtain a smooth, continuous representation of the sampled membrane structures, we interpolated the membrane heights *z*_*_ across a dense grid in the xy-plane via Gaussian Process Regression (GPR). Figures 1E, F and Fig. S6 show representative, interpolated height-profiles of the cytoplasmic leaflet. Domains of strong protrusion (dark blue) appear in the proximity of clusters composed of CAV1-8S complexes, whereas sinks (light color) occur at CAV1-depleted regions. In this way, the CG simulation displays the characteristic behavior of CAV1 sensing or inducing membrane invaginations. We quantified this relationship by computing CAV1-densities (unitless sum of Gaussian kernels) for all points of the GPR-derived membrane surfaces (Fig. 1G, S7). This approach reveals a nonlinear trend: For small CAV1 densities, we obtain a strong positive correlation to the elevation of the membrane. Proximity to a single CAV1-8S complex thus causes membrane protrusion. The correlation attenuates for larger CAV1 densities up to ~1.0, followed by a sudden increase. Larger invaginations (> 100Å, Fig 1G) hence appear at CAV1-densities of at least ~1.25, showing that the action of more than one complex is required.

By overcoming major limitations of previous computational approaches regarding length-scale and accuracy, our results hence support the idea that membrane remodeling during caveolae assembly is driven by the collective action of several caveolin-complexes. This view is in line with theoretical models and atomistic simulations.^30, 34^ We further stress this based on mean curvature profiles computed via derivatives of *z*_*_ w.r.t. *x*_*_, *y*_*_ (see Materials and Methods section). Note that typical procedures to quantify membrane bending rely on discretization techniques, whereas our GPR-based protocol facilitates more granular profiles (Fig. 1 H&I, S8). The interpolated mean curvature profiles exhibit distinguished contour lines of positive mean curvature (purple and orange dots in Fig. 1 H,I) covering CAV1-rich domains.

We highlight as well that the computed curvature profiles hence describe convex invaginations, i.e., bending into the cytoplasm by the CAV1-8S complexes, that are fully consistent with the process of caveolae budding. As illustrated in Fig. 1 H, I and Fig. S8 these invaginations can stretch over more than 50nm which is in the range of experimentally characterized caveolae assemblies^4, 35^ and reveal long-ranged interactions between CAV1-rich domains. Furthermore, these invaginations are highly dynamic as they adjust to the diffusion of the CAV1-8S complexes. This dynamics, however, is not merely stochastic but shows a pronounced preference for coordination between different complexes (Fig 1D) that is correlated to the bending of the membrane, a relationship that we further examine in the following section.

### Mechanism behind membrane remodeling by CAV1-8S complexes

To elucidate the mechanistic basis of collective membrane remodeling mediated by CAV1-8S complexes, we tracked the interactions between distinct complexes throughout the simulation. As shown in Fig 2A, diffusion can drive distant complexes into proximity that allows them to bind to each other. Intriguingly, the bound state between two complexes shows a high degree of stability and is characterized by an edge-to-edge arrangement (as the center-of-mass distances correspond to twice the radius of the CAV1-disks, ~14nm).

**Fig. 2.**
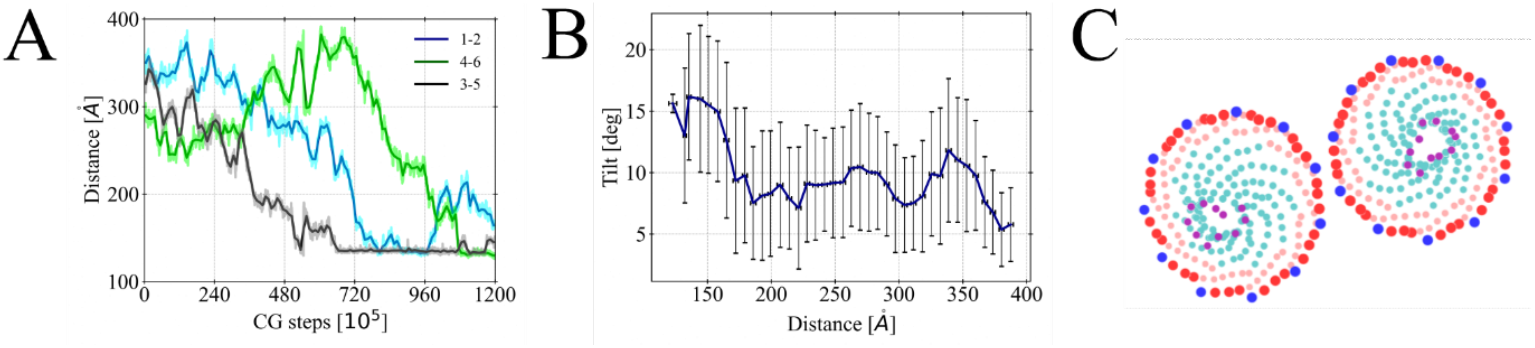
Binding of CAV1-8S complexes. **(A)** Distances between selected complexes reveal significant diffusion and transient binding. **(B)** Binding of two complexes significantly affects their orientation (tilt). Tilting is computed as angle between vectors that describe the extension of the beta-barrel (orthogonal to the flat disk). Binding of two complexes induces a mechanical coupling that is characterized by substantial tilting between the complexes. **(C)** Representative snapshot of two bound CAV1-8S complexes in CG representation. On the outer rims, negatively charged beads are highlighted in blue, and positively charged beads in red.

Furthermore, we have observed that the complexes are not rigidly embedded in the xy-plane but undergo orientational fluctuations hereafter denoted as tilting that quantifies the angle between the z-axis and direction of the beta-barrel (Fig. S9). In this way, the complexes adjust to their environment (e.g., membrane undulations), a mechanism that is particularly leveraged when two complexes approach each other: Upon binding, the convex tilting between complexes increases significantly (Fig. 2B, distances around ~15nm). Such mechanistic coupling means the emergence of a curved domain in the membrane, as the complexes are no longer in planar orientation.

Based on our CG MD simulations, we argue that the binding of different complexes is promoted by the electrostatic profile of the CAV1-8S complex. We noticed a nuanced distribution of charges characterized by an alternating 3-to-1 composition of positive and negative charges. As a consequence, distinct complexes can rotationally align to form electrostatically attractive interactions (Fig. 2C). This CG charge distribution can be projected back to the atomistic structure, so that we can identify four amino acids attributable to the binding of two CAV1-8S complexes: D82, K86, K96 and R101. We emphasize that D82, K86 and R101 show an exposed configuration (in the atomistic structure, Fig. S10) that may allow for contacts by other protein complexes. Interestingly, these residues are part of the scaffolding domain of caveolin-1, which is disease-associated and involved in numerous signaling pathways.^28, 33^ For these reasons, the scaffolding domain is hypothesized to directly mediate protein-protein interactions - an idea supported by our simulations and consistent with single molecule localization microscopy data. ^15, 33^

These results imply that the sequence of caveolin-1 may have been functionally optimized not only to encode folding into the 8S oligomers, but also higher-order interactions essential for caveolae formation. Figures 3A, B depict representative simulation snapshots illustrating how the formation of such higher-order interactions forms interplay with the membrane-structure: Isolated CAV1-8S complexes (Fig 3A) bend the exoplasmic leaflet locally at the center of the complexes (due to hydrophobic interactions with the beta-barrel).^30^ Upon approaching each other, the stress profiles of two complexes merge (Fig 3B) which leads to convex tilting and in this way to local elevation of the membrane domain (Figs 3B, C).

**Fig. 3.**
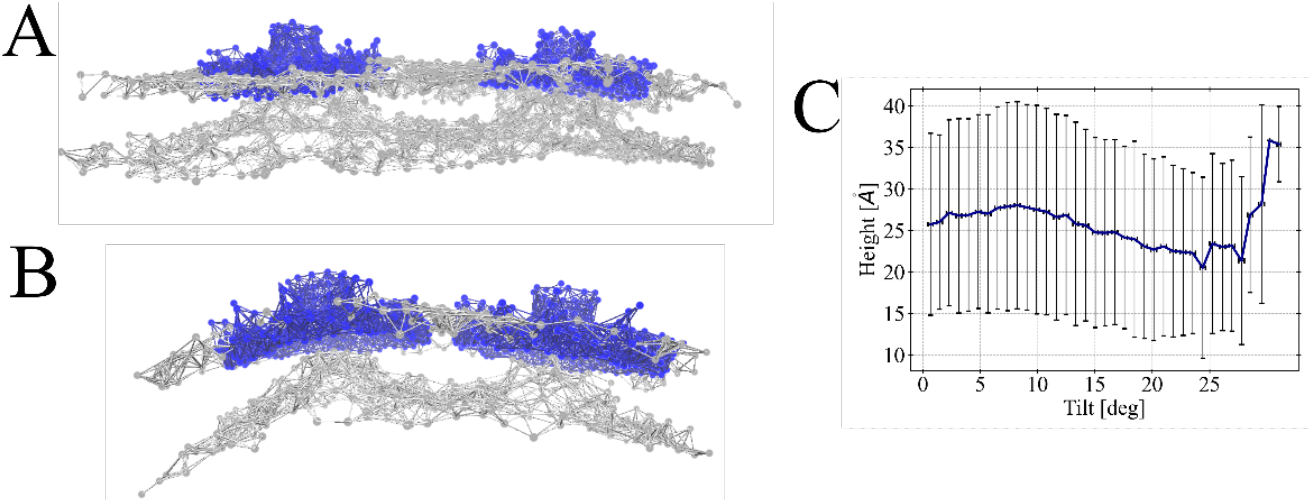
Coupling between CAV1-8S binding and membrane remodeling. **(A, B)** Representative snapshots in separated and bound form (blue: CAV1-8S complexes, silver: lipid bilayer). In the former case, the complexes bend the exoplasmic leaflet locally under the beta barrel, whereas binding leads to a superposition of the deformation profiles in the exoplasmic leaflet. **(C)** Larger membrane heights preferentially emerge when complexes are strongly tilted which quantifies the mechanical coupling between orientation of the CAV1-8S complexes and protrusion of the local domain.

### Formation of large-scale invaginations

While the aforementioned simulation has given fundamental new insight into the collective functioning of caveolin-1, the methodology is slightly limited by two factors: First, the constant box size may suppress stronger membrane curvature. Second, simulating transition from isolated systems to an organized cluster is an inherent sampling issue even for CG modeling. The insights discussed above, however, allow us to design a simulation-setup that addresses these issues.

Note that the binding pose shown in Fig 2C occupies approximately two of the eleven edges of one CAV1-8S complex. Thus, we suggest that one complex can be coordinated by six others, and we then started simulations from a structure arranged in this way (Fig 4A). During that simulation, we continuously compressed x- and y-dimensions to screen the energy landscape for membrane curvature generation and hence to sample formation of ~70nm large membrane invaginations scaffolded by the CAV1-cluster (Fig. S11, Movie S3&S4). This setup can be seen as a computational analog to experiments (e.g., optical stretching or magnetic tweezers) that probe the buffering of membrane tension by caveolae assemblies.^20, 36^

**Fig. 4.**
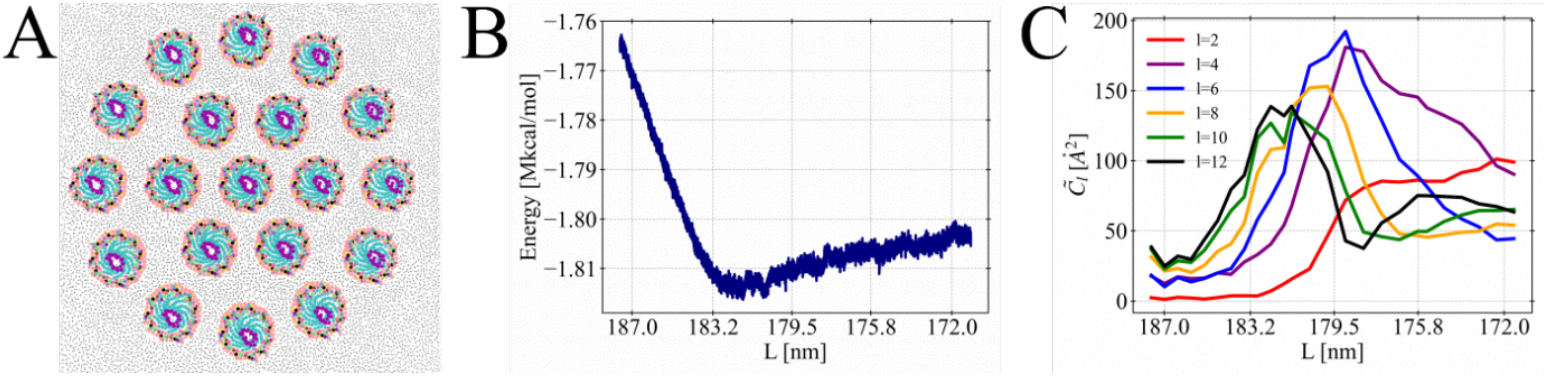
Box compression simulations lead to large-scale invaginations mediated by a CAV1-cluster. **(A)** Top view of the starting structure (zoomed in on the central membrane domain that is covered by 19 CAV1-8S complexes). The beta-barrels are highlighted in purple. **(B)** Recorded energy landscape during compression of the box, with a minimum around ~181nm. **(C)** Symmetric modes during simulation inferred from a spherical harmonic analysis of the central membrane invagination covered by the complexes.

The screened energy profile shows a minimum at intermediate curvature, but the system remains relatively flexible with respect to further compression/bending (Fig. 4B, S12). While we do not observe significant readjustment in POPC/CHOL ratios (Fig. S13), the shape of the invagination undergoes distinct changes as quantified with a spherical harmonic analysis (see Methods). Initial budding shows systematic increases in the symmetric coefficients (Fig. 4C), after reaching the energy-minimum, however, the symmetric modes begin to attenuate, and small or asymmetric modes dominate (Fig 4C, S14). We note that cryo-electron tomography experiments have revealed polygonal shapes for the caveolae vesicles.^4, 14, 37^ Since the minimum of the energy profile coincides with the maximum of the l=12 mode (Fig. 4 B, C), our CG simulations suggest that CAV1-8S complexes collectively scaffold the membrane to deformations with a noticeable polyhedral symmetry. Additional cofactors may then be needed to preserve the symmetry for further restriction of the invagination.

The CG simulation also substantiates our previously discussed mechanistic coupling between membrane bending and tilting of the CAV1-8S complexes (Fig. 5). Formation of the ~70nm large invagination correlates with increased tilting of the complexes, especially for those at the contour of the invagination that can tilt by more than 50° (Fig. S15). Thus, our simulation shows the formation of a caveolae-bud with very similar characteristics to hypothesized models for caveolae vesicles that suggest that CAV1-8S complexes are embedded in flat, local membrane surfaces of different orientations forming a polyhedral shape.^4, 27^ Note that the formation of CAV1-mediated membrane invaginations was also confirmed in an additional simulation of five complexes, where we have observed clustering of three complexes that localizes the membrane stress upon compression (Fig. S16-S18).

**Fig. 5.**
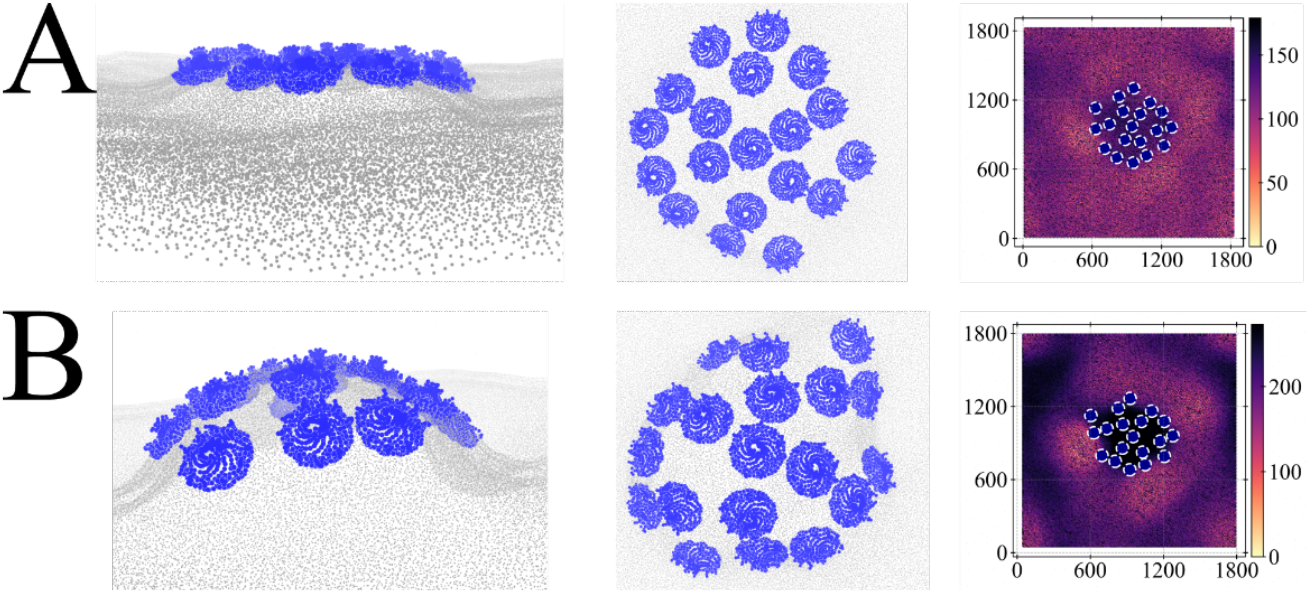
Structures of the formed invaginations. **(A)** Top row shows structure of the invagination at the energy minimum in side (left) and top view (middle). The z-profile (in Å units, right) interpolated via GPR quantifies formation of an invagination at the CAV1-cluster. **(B)** Recorded representative structure for stronger compression (*L*~174*nm*) shows pronounced budding. Complexes at the invagination boundary exhibit substantial tilting (middle). Increased levels of membrane elevation are evaluated with GPR (right).

## Discussion

Caveolin-1 proteins mediate substantial remodeling of the membrane which can be critical to many cellular processes. The mechanistic basis for their functioning has remained elusive, since experiments offer only limited resolution, and all-atom MD simulations only limited length-scales. Previous atomistic simulation studies have therefore largely focused on simulations of a single CAV1-8S complex embedded in one leaflet of a lipid bilayer. These simulations show only very localized deformations of the membrane shape,^29, 30^ hence suggesting that the flat-to-curved transition in caveolae biogenesis must be orchestrated collectively by numerous complexes.

Coarse-grained simulations, on the other hand, have the potential to provide for both high resolution and scalability if done as rigorously as possible. Top-down CG models, however, appear to cause membrane curvature at odds with all-atom simulations for such a monotopic system. For this reason, we have trained a bottom-up CG model that captures the insertion of the CAV1-8S complex as well as relevant membrane properties. To the best of our knowledge, this is the first bottom-up model for such a system. We therefore emphasize that our protocol may be extended to a broader class of protein-membrane systems, e.g., peripheral or transmembrane proteins.

In our CG MD simulations, we have observed significant diffusion of the individual CAV1-8S complexes with a strong tendency for clustering. Leveraging Gaussian Process Regression to densely interpolate the membrane surface, we find a nonlinear correlation between CAV1-occupancy and membrane height, that can result in ~ 50nm broad membrane invaginations. In that case, collective action between the complexes is also enhanced by electrostatic interactions, as we noted that in the correct rotational alignment the complexes may attract each other. This attraction can be attributed to four amino acids in the scaffolding domain, D82, K86, K96 and R101. Attractive electrostatic interactions between scaffolding domains of two 8S complexes is a novel, testable hypothesis. We argue that these interactions are structurally very plausible, as the charged amino acids are in place to form salt bridges. It is also fully in line with network analysis of single molecule localization microscopy and biochemical fractionation experiments that captured dimerization of 8S complexes (referred to as S1B scaffolds). ^14–16, 38^ Dimers of CAV1-8S then likely drive lattice formation into the polyhedral 70S complex. Strikingly, the F92A and V94A mutations in the scaffold domain have been shown to maintain competence of caveolae assembly but have a pronounced impact on the size of the scaffolds (including S1B)^15^. In addition to these hydrophobic mutants, mutation of R101 has also been related to disease.^28^

In addition, our CG simulations suggest that proximate complexes respond to each other’s stress profile in the exoplasmic leaflet by convex tilting. We emphasize that this represents a direct biophysical explanation and partial structural evidence of the dynamic curvature mechanism proposed by Lim and coworkers.^33^ Such dynamic curvature generation facilitates access by further cofactors and has therefore been hypothesized to modulate signaling pathways such as eNOS.^33^

Compressing the lateral-dimensions of the membrane, triggers these mechanistic couplings and therefore forms ~70nm large invaginations at the center of the CAV1-cluster. We find an energy minimum for intermediate invaginations that are characterized by maxima in the high l-modes obtained from a spherical harmonics analysis. While the energetic cost to further restriction is reduced, it suggests that additional cofactors are needed to further restrict the invagination and maintain the polyhedral symmetry. Thus, our CG simulations show that assembled CAV1-8S complexes self-organize to scaffold initial membrane invaginations – in agreement with high-resolution imaging techniques and fragility assays.^14, 16, 20^

In the future, it will be of value to include cavin-trimers and PIP2 lipids (that bind to the HR1 domain of cavin^39, 40^) in our CG model. Such simulations are of particular interest and are likely to extend the biophysical and molecular view of caveolae assembly from this paper, as cofactors may also sense the electrostatic profile of the CAV1-8S complex.

## Materials and Methods

### MD simulations

Atomistic MD simulations were performed with the Gromacs/2020 package for two different systems, a CAV-1 8S complex embedded in a lipid membrane with 70/30% POPC/CHOL composition, and a pure membrane system of same composition.^41^ Resulting trajectories were used to train the CG models. For the CAV-1 system, we used the output structure from a previously published Metadynamics simulation as starting structure.^30^ This system was simulated in the constant NVT-ensemble without any bias for ~ 80ns, using otherwise the same simulation-setting as in the Metadynamics simulation. The membrane system was generated with CHARMM-GUI adjusting x- and y-dimensions of 40nm and an extension in z-direction of 11nm.^42^ The Charmm36m force field was employed to describe interactions between the atoms of the systems.^43^ Energy minimization was carried out for 5,000 steps using the steepest descent method. The membrane system was then equilibrated following the CHARMM-GUI generated input-files:^44^ First, the system was simulated at a reference temperature of 310 K employing a Berendsen thermostat with a coupling constant of 1ps.^45^ This equilibration was performed in the NVT-ensemble for 250,000 time steps. Subsequently, a semi-isotropic Berendsen barostat was applied with a reference temperature of 1bar, a compressibility of 4.5 ∙ 10^-5^ *bar*^-1^ and a coupling constant of 5 ps.^45^ The first 375,000 steps of the entire equilibration were performed with a time-step of 1fs, and the remaining 750,000 steps with 2fs. Through equilibration, conventional restraints were used and gradually reduced. The production run was performed for ~50ns in the NVT-ensemble using a Nose-Hoover thermostat.^46, 47^

### Coarse-grained modeling

Mapping rules for the lipid and caveolin-1 molecules were derived based on the essential-dynamics coarse-graining (ED-CG) method using the OpenMSCG package.^48, 49^ Each cholesterol and POPC lipid molecule was mapped to three and six CG beads, respectively (Fig S1). For the CAV1-8S complex, identical mapping was performed for each monomer, resulting in 30 CG beads per caveolin-1, and hence 330 beads per CAV1-8S complex (Fig S1). This corresponds to a resolution of 4-5 amino acids per bead. The CAV1-8S disk was then modeled as a hetero-elastic network model (hENM) parameterized with OpenMSCG using a cutoff of 30 Angstrom.^48, 50^ The CG potentials for bonds and angles of the lipids were derived with force matching as tabulated (4^th^-order B-spline interpolated) potentials.^51, 52^

Non-bonded interactions were trained iteratively through the relative entropy minimization (REM) technique.^53^ In the first step, we trained non-bonded lipid-lipid and lipid-protein interactions as tabulated 4^th^-order B-spline interactions. To increase stability of the protein-lipid interactions, we have replaced the tabulated protein-lipid interactions by training truncated Gaussian potentials in the second step of training. In consecutive steps, we have trained perturbations in the form of 9-6 Lennard-Jones and truncated Gaussians. The objectives for these perturbations are to make the CG-model transferable also to a pure membrane system and to more accurately capture the bending modulus. Additionally, excluded volume interactions were added between protein and lipid beads to increase repulsion in the marginally sampled short-range distances. A detailed overview of the training is given in Table S1 of the SI. As shown in Fig. S2 and Movie S1, REM training of the protein-membrane interactions results in a correct monotopic binding mode of the CAV1-complex, and our procedure also facilitates self-assembly of the lipid membrane (Movie S2) and captures the bending modulus in quantitative agreement with experiments (*κ*_*b*_ = 29.6 *k*_*B*_*T*, Fig. S3). Interactions between different CAV1-8S complexes were described as Debye-screened and excluded volume interactions. The functional form of the full model hence is given by:

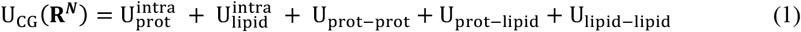

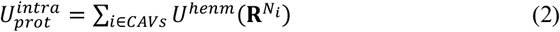

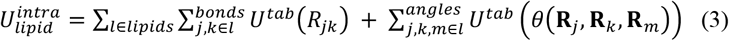

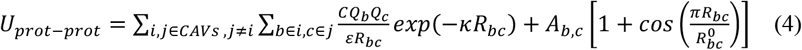

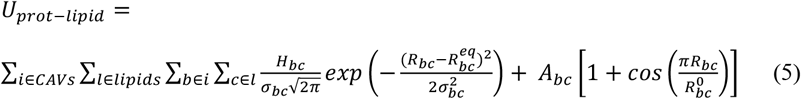

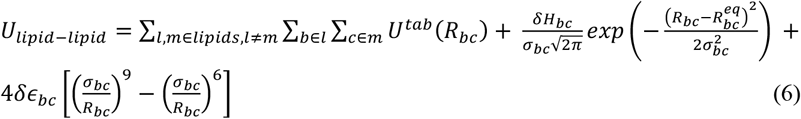

For the screened electrostatic interactions between CAV1-8S complexes, we used *ε* = 2 and *κ* = 0.12*i*4Å^-1^. C converts to units of energy. Following previous studies^54, 55^, the charges of the protein beads *Q*_*b*_ and *Q*_*C*_ were assigned as sum of the partial charges of all atoms mapping to the corresponding CGbead. For protein-protein interactions, the repulsion coefficient *A*_*b,C*_ were set in a range from 0 to 18.75 kcal/mol, depending on charges and radial distance from the center of the complex. Analogously, 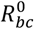 was set in a range from 0.0 to 12.0Å. For protein-lipid interactions, *H*_*bC*_ was trained via REM based on σ_*bC*_ = 2.0Å and 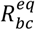 as mean-distance (measured in the atomistic reference simulation), with exclusions specified by *A*_*bC*_ = 25 *kcal*/*mol* and 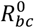 at the transition between repulsion and attraction. Refinements of the lipid-lipid interactions were trained with a σ_*bC*_ corresponding to the width and 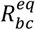 to the position of the first peak in the radial-distribution function, constraining |*δH*_*bC*_| < 0.15 *kcal*/*mol* Å and |*δϵ*_*bC*_| < 2 ∙ 10^-4^*kcal*/*mol*. Apart from the hENM-interactions, a cutoff of 25.0 Å has been used in all CG simulations.

### CG system initialization and simulation

All CG simulations were performed with the LAMMPS simulation package.^56^ The topology-files that also contain the starting configuration were built with in-house developed python-scripts. Following recent suggestions,^57^ REM-training was started from disordered structures which were generated with packmol.^58^

Large-scale CG simulations were performed for three different systems with different numbers of CAV1-8S complexes: 12, 19 and 5 (termed 12CAV, 19CAV and 5CAV in the following). The x- and y-dimensions for the three systems were 126.4nm, 187.6nm and 113.4nm, respectively. Magnitudes of the the z-dimensions are of lesser significance, since the models are solvent-free, and were set to 200.0, 200.0 and 50.0 nm, respectively. Periodic boundary conditions were applied.

The CG systems were minimized for 1,000 steps with the conjugate gradient method and afterward simulated at T=310K using a Langevin-thermostat with a coupling time-constant of 10ps.^56^ Neighbor lists were updated every step, and all systems were simulated in the constant NVT-ensemble, except the compression-simulations. Coordinates were written out every 12,500 steps (12CAV) or 25,000 steps (19CAV, 5CAV). An integration time-step of 15fs was employed. The systems were simulated for ~120 ∙ 10^6^, 14.*n* ∙ 10^6^, 19.3 ∙ 10^6^ (12CAV, 19CAV, 5CAV) CG steps. The compression rates for the 19CAV and 5CAV systems were ~1.0 *and* 0.3 *nm*/(10^6^ *CG steps*), respectively. We performed three additional replicates for the 12CAV system starting from different velocities for 65 ∙ 10^6^ CG steps (Fig. S5, S7). Similarly, two additional compression-simulations were performed for the 19CAV systems (Fig. S12).

A randomized lipid structure was simulated for 14.7 ∙ 10^6^ steps in the NVT ensemble (with x- and y-dimensions of 31.6nm) at 310K. The last 0.7 ∙ 10^6^ steps (after reassembly) were used to estimate the bending modulus.

The trajectories were processed with the MDAnalysis library for subsequent analysis.^59^ Normal vectors on the disks were computed as vector between center of masses of beads with type 20 and beads of type 28 (direction of the beta barrel).

### Estimation of the bending-modulus of the lipid-model

The lipid-bilayer was discretized in xy-area with a grid-spacing of 5Å, and only POPC-headgroups were considered. Following Pöhnl et al,^60^ the z-profile was computed by weighting the contributions of the beads based on their xy-distances from the bin-centers by a Gaussian function with σ = *n*Å. For both leaflets, we computed the Fourier coefficients summing over all bins N:

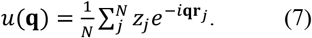

The mean Fourier coefficient over both layers was used in the interpolation of the bending modulus *κ*_*b*_ via:

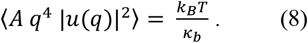

Here, the constant A denotes the membrane surface area. Interpolation was carried out in the low q-region (Fig S3), and yielded *κ*_*b*_ = 29.6 *k*_*B*_*T*.

### Gaussian Process Regression

The z-coordinates of the lipid head-groups were interpolated at a spacing of 5Å in x- and y-dimension (denoted **r**_*_ = (*x*_*_, *y*_*_)) via Gaussian process regression^61, 62^:

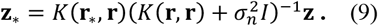

Here, **r** = (**x, y**) and **z** are the sampled POPC headgroups, and K denotes kernel matrices,

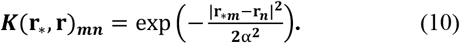

We used *α* = 1.0Å and σ_*n*_ = 1.0 ∙ 10^-4^ for interpolations of the height profiles and *α* = 2.5Å with σ_*n*_ = 0.01 for curvature analysis. **z**_*_ was regressed over 5×5 nm^2^ windows over the xy-surface, using training data of 8×8 nm^2^ windows, where the 5×5 nm^2^ windows are centered.

Mean curvature was computed by:

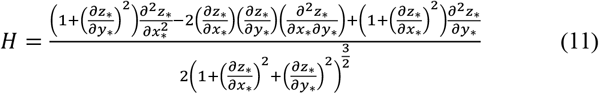

The partial derivatives are calculated straightforwardly from the derivatives of the Kernel, e.g.:

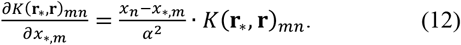

### Spherical harmonics

In the analysis of the membrane shapes, the center of the coordinate system was placed inside the membrane invaginations, and Cartesian coordinates of the POPC lipid head-groups were transformed to the spherical coordinate system. Membrane shapes were then characterized with spherical harmonics:

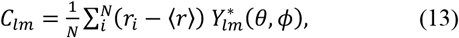

with 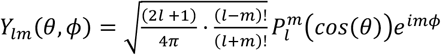 and 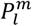 are the associated Legendre polynomials.

The membranes surfaces were then characterized by:

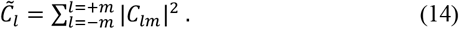

This analysis was performed for a circular domain of the exoplasmic leaflet with a radius of 45nm centered at the CAV1-cluster for every 10^th^ frame of the output trajectory.

### Caveolin density and coordination

The caveolin density was computed for every grid-point (**r**_*_):

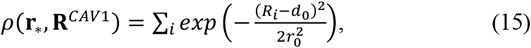

where *R*_*i*_ is the center-of-mass distance of a CAV1-8S complex (indexed i) to a grid-point. Distances were computed as 2d (x- and y-dimension), weighted by *d*_0_ = 70Å and *r*_0_ = 100Å. Cooperativity between the complexes was computed as

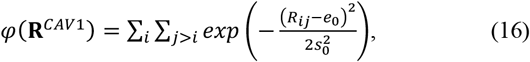

using *e*_0_ = 140Å and *s*_0_ = 50Å. The choice of *d*_0_ and *e*_0_ are motivated by the radial extension of the CAV1-8S disks. The correlations between membrane height and CAV1 density were computed for the second half of the trajectories.

## Supplementary Materials

### The PDF file includes

Figs. S1 to S18

Table S1

Legends for Movies S1 to S4

Information on the development of the CG model and replicate simulations

## Funding

This project received funding from the European Union’s Framework Programme for Research and Innovation Horizon Europe (2021–2027) under the Marie Skłodowska-Curie Grant Agreement No. 101109916 to K.L. Research reported in this publication was also supported by the National Institute of General Medical Sciences of the NIH under award number R01GM063796 to G.A.V. The content is solely the responsibility of the authors and does not necessarily represent the official views of the NIH. Computational resources were provided by the University of Chicago Research Computing Center, the NIH-funded Beagle-3 computer (NIH award 1S10OD028655), and by the Frontera supercomputer at the Texas Advanced Computing Center (TACC) at the University of Texas at Austin and funded by the National Science Foundation (NSF) (grant OAC-1818253).

## Author contributions

K.L. and G.A.V. designed the research. K.L. developed the coarse-grained model, designed and performed simulations, developed the software, analyzed the data, prepared the figures, and wrote the manuscript. G.A.V supervised the research and contributed to the writing of the manuscript.

## Data and materials availability

The coarse-grained model is freely available at 10.5281/zenodo.16105882. This deposit also includes sample input files and scripts to generate and analyze large-scale simulation systems.

## Notes

### Competing Interest Statement

The authors have declared no competing interest.

