## Supplementary Materials for "Membrane Remodeling by the Collective Action of Caveolin-1"

This file provides further information on the training of the CG-models, replicate simulations, orientational fluctuations and membrane remodeling by Caveolin-1.

### 1. Development of the coarse-grained (CG) model

Each of the eleven protomers of the CAV1-8S complex is mapped to 30 beads (Fig. S1 A), applying identical mapping for the protomers. Cholesterol is mapped to three beads, and POPC to six beads.

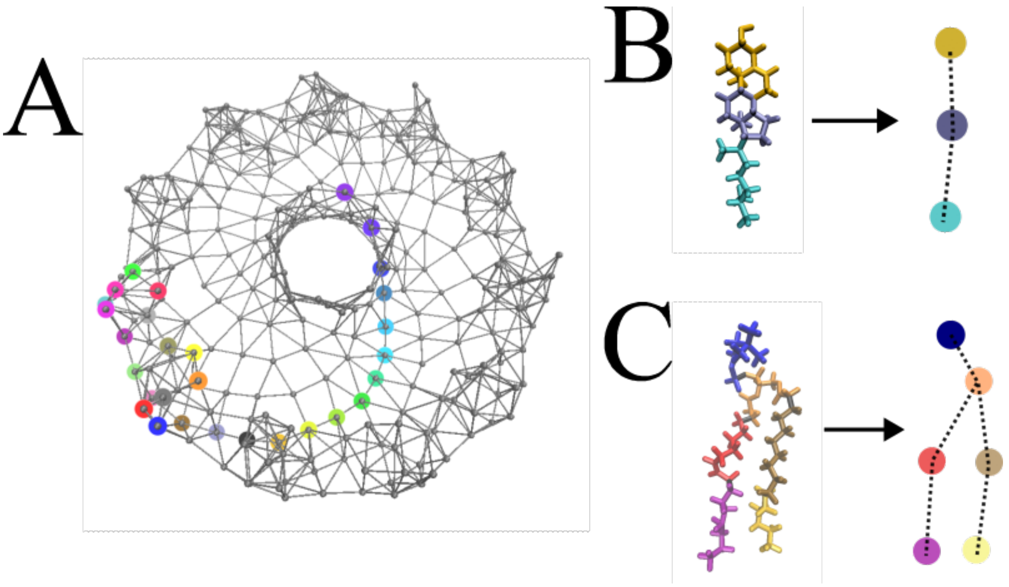


**Fig. S1. Coarse-grained representations of the system. (A)** Protomers of the CAV1-8S complex are mapped to 30 beads, the representation of one protomer is highlighted by colored beads. The vertices schematically represent the elastic network model. **(B)** Mapping of Cholesterol, dashed lines indicate bonded interactions in the CG-model. **(C)** Mapping of POPC, dashed lines indicated bonded interactions.

Iterative optimization of non-bonded lipid-lipid and protein-lipid interactions via REM (eq. S1) allows us to learn the monotopic embedding of the CAV1-8S complex (Fig. S2).

In REM, we perform CG interactions for each optimization to update the parameters of the CG force field according to:

$$\lambda_{i+1} = \lambda_i - \sigma \frac{\left(\langle\frac{\partial U}{\partial \lambda}\rangle_{AA} - \langle\frac{\partial U}{\partial \lambda}\rangle_{CG}\right)}{\beta\left(\langle\left(\frac{\partial U}{\partial \lambda}\right)^2\rangle_{CG} - \langle\frac{\partial U}{\partial \lambda}\rangle_{CG}^2\right)}$$

The entire training process consists of 5 steps (Tab S1). Step 1 & 2 are the main steps. In step 1, tabulated lipid-lipid and protein-lipid interactions are trained. This sets the membrane in place and configures the CAV1-8S complex appropriately. However, these interactions are prone to misconfigurations between the protein complex and the lipids on longer timescales. Thus, we discarded the tabulated protein-lipid interactions, and retrained them as Gauss-cut potentials (step 2, Fig. S2, Movie S1). The remaining three steps are refinements to the model. 9-6 LJ parameters are trained to allow for larger membrane undulations and hence a lower, and more realistic bending modulus (Fig. S3). Note that realistic description of the bending modulus of lipid bilayers has been a challenge for solvent-free, bottom-up CG models.[1] In step 4, minor parameter modifications were trained to enhance transferability and robustness. To avoid overfitting of interactions[2] between cholesterol and beta barrel beads, in step 5, we rescaled and retrained those interactions so that the beta barrel channels ~2-3 cholesterol molecules (in consistency with atomistic simulations).

Importantly, our lipid model also qualifies for correct self-assembly starting from a randomized structure (Movie S2), which demonstrates robustness of the model also on longer timescales. Furthermore, radial distribution functions obtained with our CG model show good agreement with the atomistic reference (Fig. S4).

**Tab. S1. REM-training of the CG-model.** First two stages train the core-of the model. Stages three and four train perturbations (denoted by $\Delta$) to train specific features. In stage five, interactions between cholesterol and the C-terminal bead-types are retrained. The column specifying the number of simulation steps (5$^{th}$ column) is in multiples of $10^6$.

| System | interactions | type | objective | steps | software | Learning-rate | Iterations |
|---|---|---|---|---|---|---|---|
| CAV1/lipid | Lipid-lipid<br>Protein-lipid | tabulated | Initial | 2.1<br>(1.5) | openmscg | 0.002 | 141 |

| | | | | | | | |
|---|---|---|---|---|---|---|---|
| CAV1/lipid | Protein-lipid | Gauss | Robustness | 1.08 (1.0) | In-house | 0.05 | 40 |
| lipid | Lipid-lipid | LJ (Δ) | Bending modulus | 0.6 (0.56) | In-house | 0.02 | 50 |
| CAV1/lipid | Protein-lipid Lipid-lipid | Gauss (Δ) | Transferability, Robustness | 1.41 (1.25) | In-house | 0.05 | 14 |
| CAV1/lipid | Barrel-Chol | Gauss | Cholesterol channeling | 1.41 (1.25) | In-house | 0.01 | 9 |

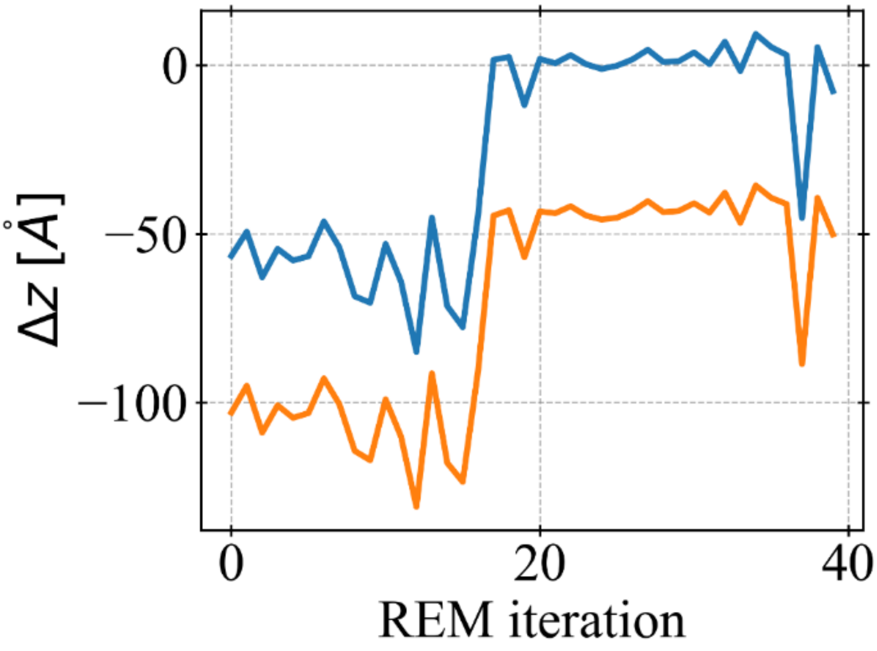


**Fig. S2. REM-training leads to monotopic configuration.** The blue line shows distances in z-direction between the CAV1-8S disk (beads of type 12), and the POPC beads of the cytoplasmic layer. Values of 0 indicate embedding in this leaflet, and distances to the exoplasmic layer are shown in orange. This plot shows the second training from Tab S1.

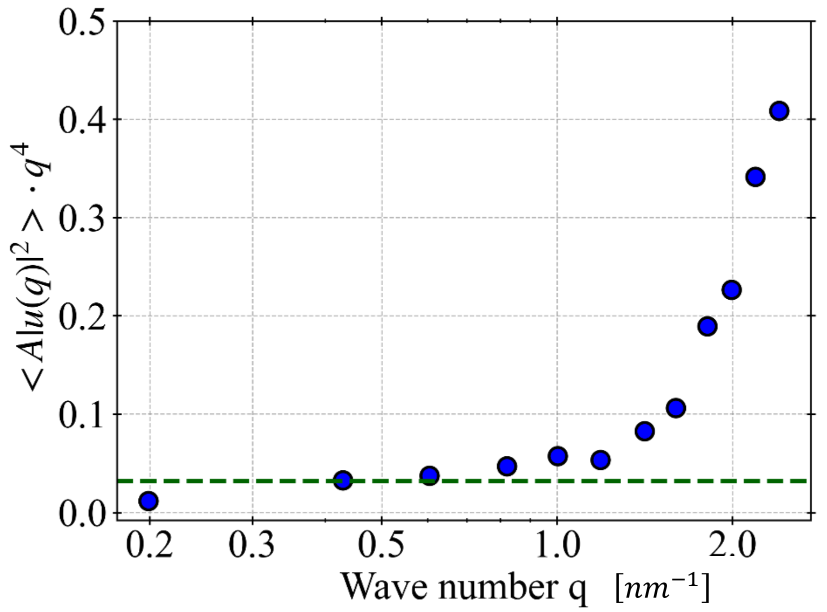


**Fig. S3. Undulation spectrum of the trained lipid bilayer.** The dashed green line is the fit in the low q-region. The bending modulus is obtained as reciprocal value of the fit.

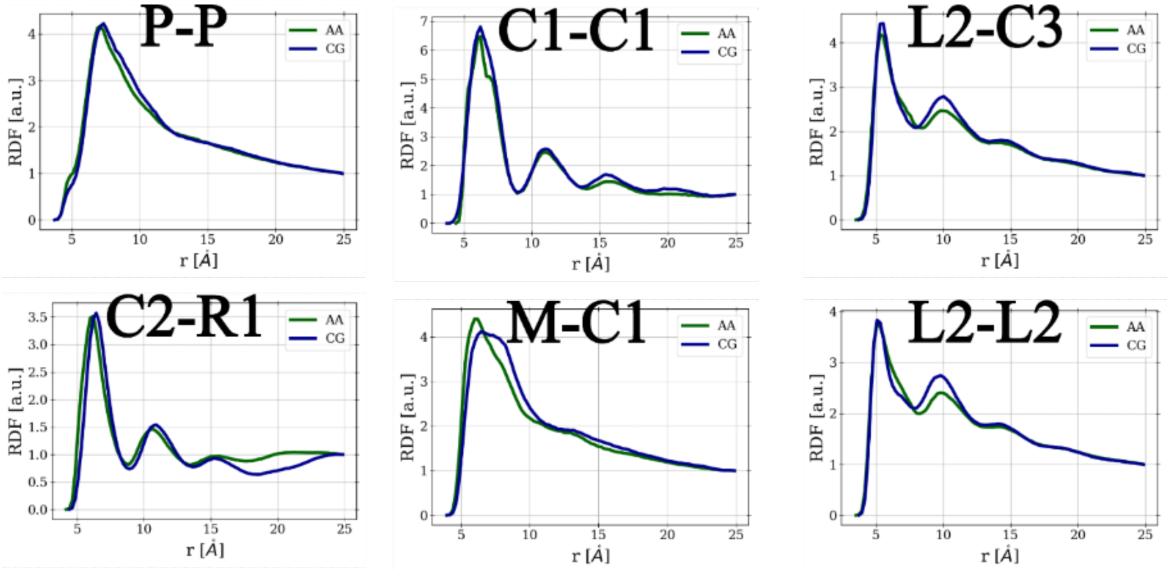


**Fig. S4. Radial distribution functions obtained from atomistic (green) and coarse-grained (blue) simulations for the lipid bilayer**. P, M, R1, L2 denote head-, middle-, and tail-groups of POPC. C1, C2, C3 denote the hydroxyl-end-, middle- and end-group of cholesterol.

### 2. Membrane remodeling and orientational fluctuations in the unbiased system

This section provides further results obtained for the simulation system that includes 12 CAV1-8S complexes. It also includes results from three replicate simulations starting from different velocities (Fig. S5, S7). Height- and curvature-profiles sampled during the simulation discussed in the main text are shown in Fig. S6 and Fig. S8.

Tilting of the complexes with respect to the z-axis are plotted in Fig. S9.

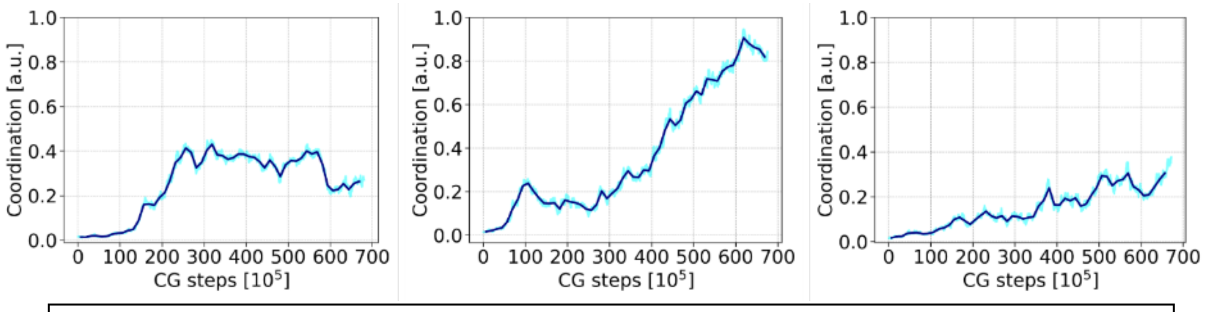


**Fig. S5. Coordination of the CAV1-8S complexes during the three replicate simulations.** The replicate simulations show dynamic binding- and unbinding-dynamics with a tendency to cluster formation similarly to Fig. 1D of the main text.

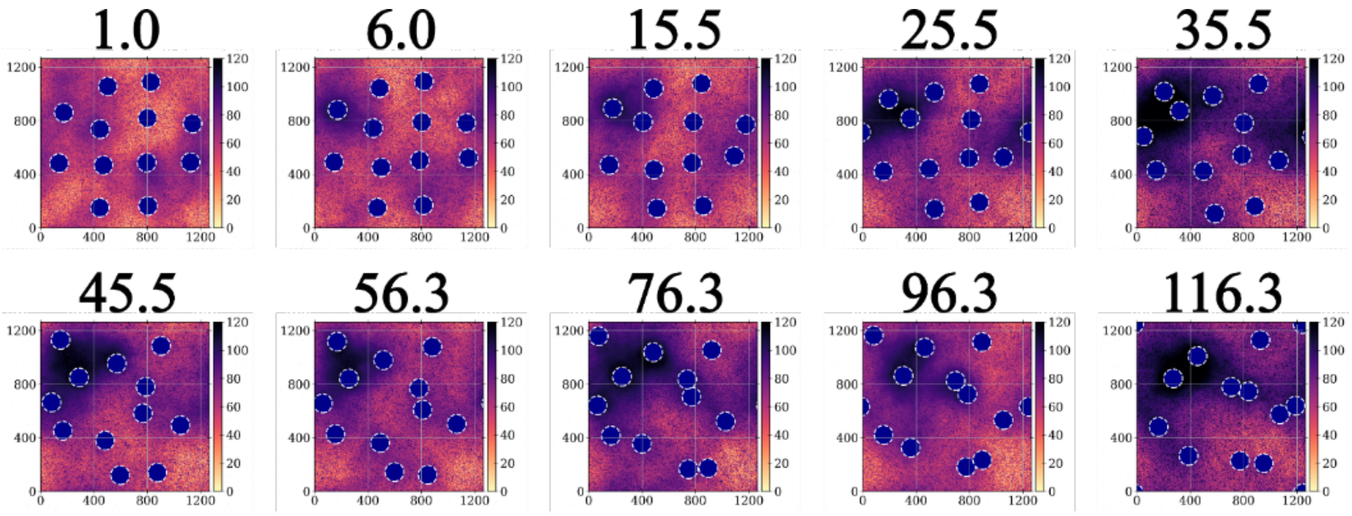


**Fig. S6. Height profiles computed via Gaussian Process Regression**. Numbers on top of the profiles specify the timestamp of the simulation in units of 1mio.

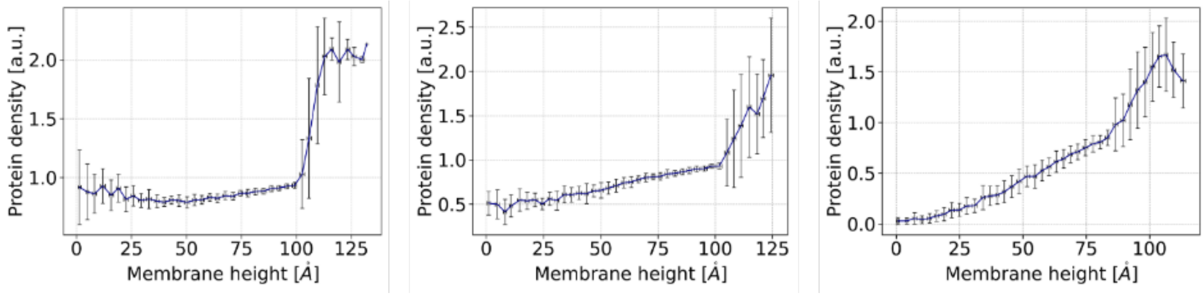


**Fig. S7. Correlation between membrane height and protein density obtained from the three replicate simulations.** The positive correlation and non-linear trend characterized by a sharp increase for protein-densities around ~1.0 is fully consistent with the discussion in the main text.

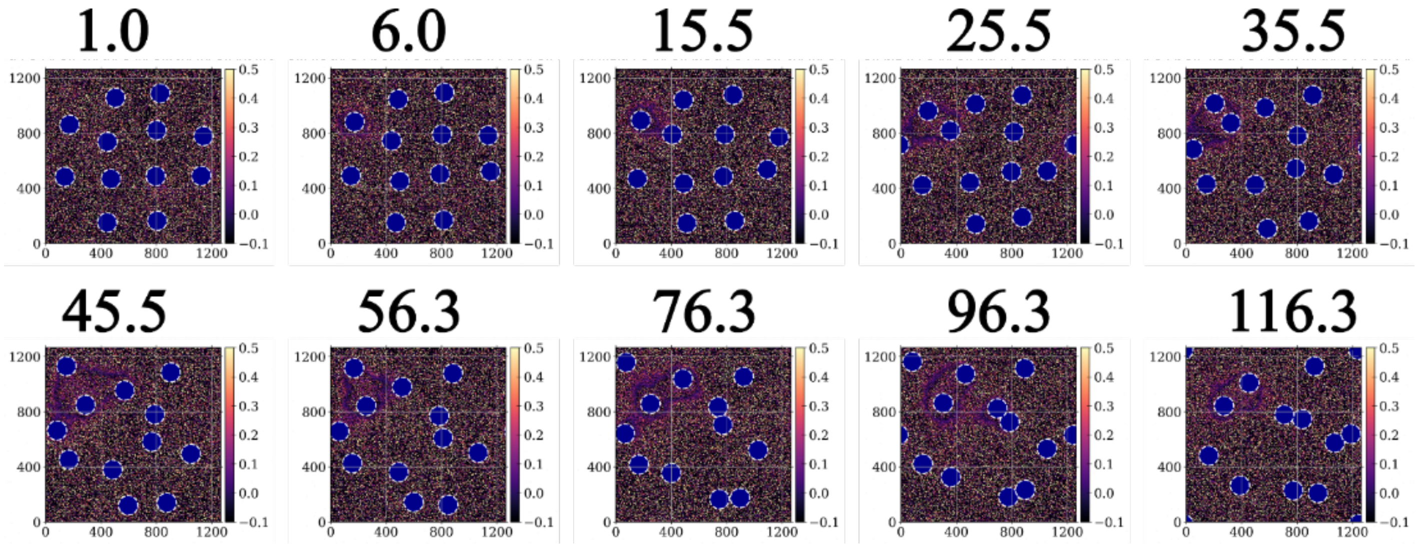


**Fig. S8. Mean curvature profiles computed via Gaussian Process Regression**. Numbers on top of the profiles specify the timestamp of the simulation in units of 1mio.

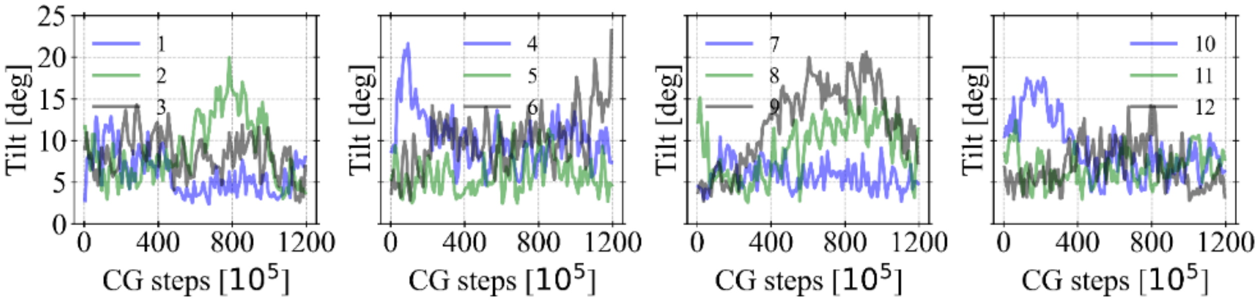


**Fig. S9. Orientational fluctuations of the CAV1-8S complex.** Tilting of the complexes (angle between extension-vector of the beta-barrel and the z-axis) shows strong fluctuations during simulation.

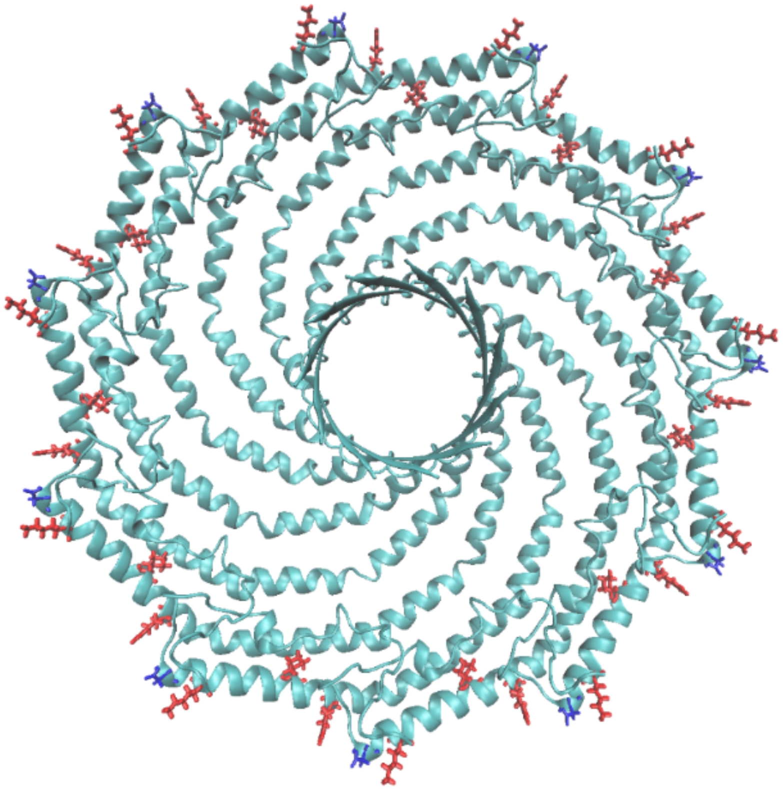

**Fig. S10. Secondary structure of the CAV1-8S complex.** Charged amino acids of the outer rims are highlighted, ASP82 in blue, and LYS86, LYS96 and ARG101 in red.

### 3. Membrane remodeling and orientational fluctuations in box-compressing simulations

This section shows results obtained from compressing the simulation box for the system with 19 circularly arranged CAV1-8S complexes (Fig. S11-15). Note that two replicate simulations (Fig. S12) stress the confidence of the simulation results. In addition, the characteristic behavior discussed in the main text was also obtained from a simulation with 5 CAV1-8S complexes. This system was also simulated using a smaller compression rate and placing the center of the cluster outside the box center to check the sensitivity with respect to the simulation settings.

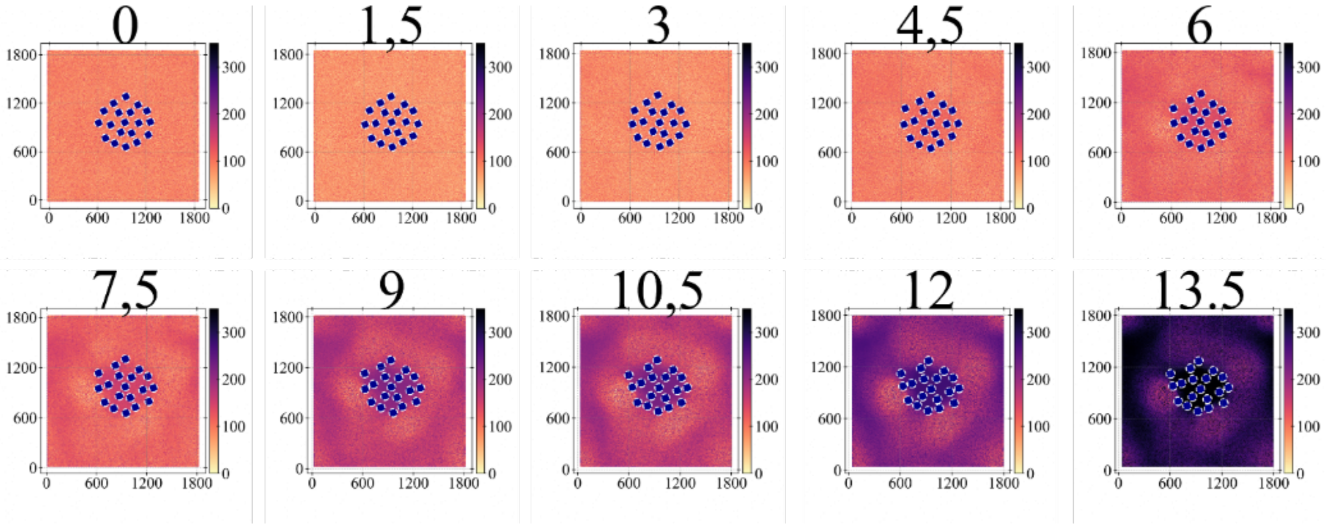


**Fig. S11. Height profiles for the system with 19 CAV1-8S complexes under compression computed with Gaussian Process Regression**. Numbers on top of the profiles specify the timestamp of the simulation in units of 1mio.

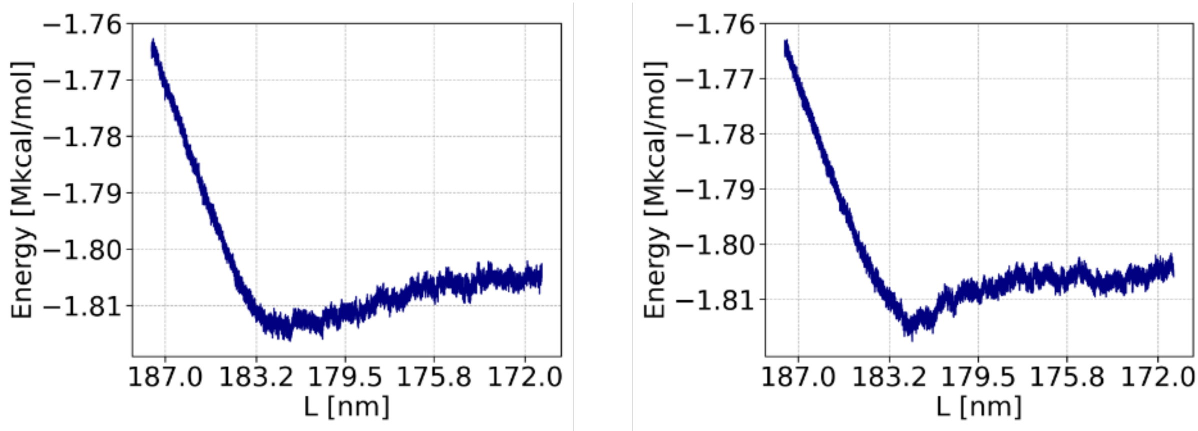


**Fig. S12. Energy profiles for replicate compression simulations of the 19 CAV1-8S complexes**. Replicate simulations (starting from different velocities) reproduce the energy profile shown in Fig. 4B.

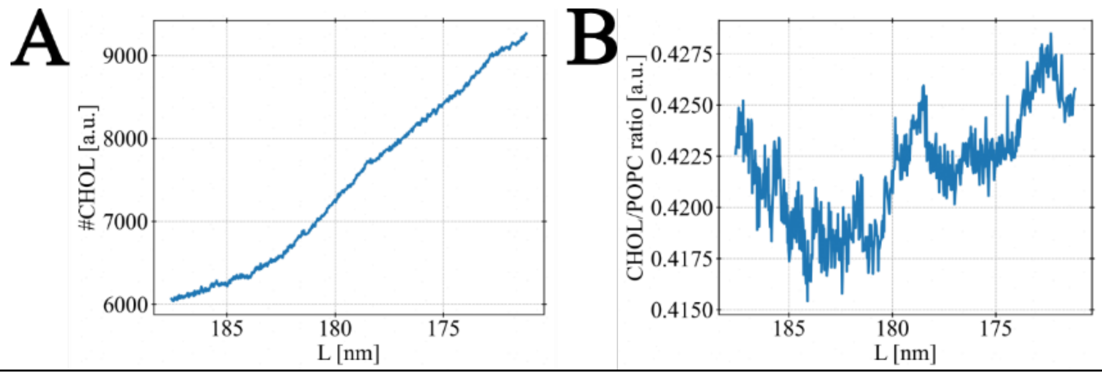


**Fig. S13. Lipid enrichment in the invagination process formed in the presence of 19 CAV1-8S complexes. (A)** Number of cholesterols increases as the invagination becomes larger. **(B)** Fluctuations of the ratio between cholesterol and POPC do not show an equivocal trend and are overall small. The profiles were calculated over the local CAV1-cluster domain where the invagination forms (analogously to the spherical harmonic analysis).

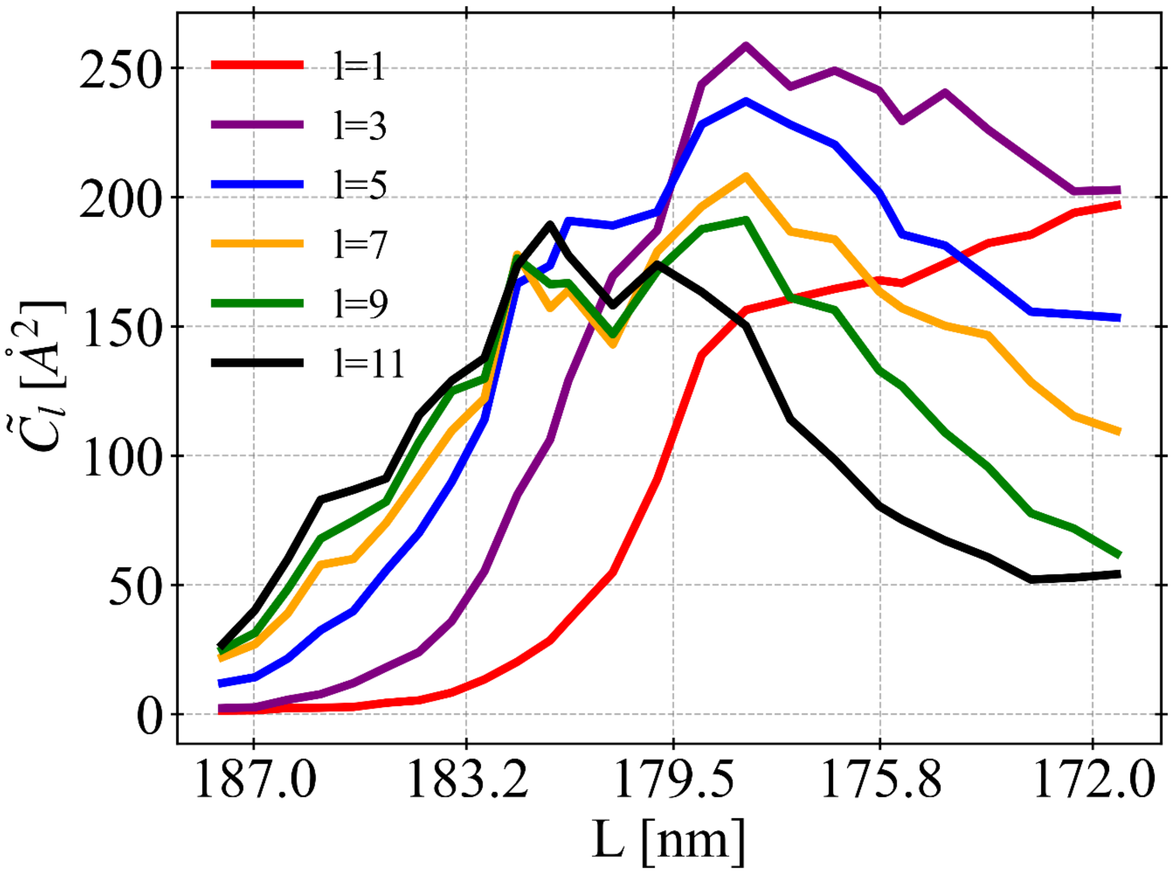


**Fig. S14. Odd spherical harmonics coefficients computed during compression of the system with 19 CAV1-8S complexes.** The coefficients were computed as described in the Materials and Methods section of the main text.

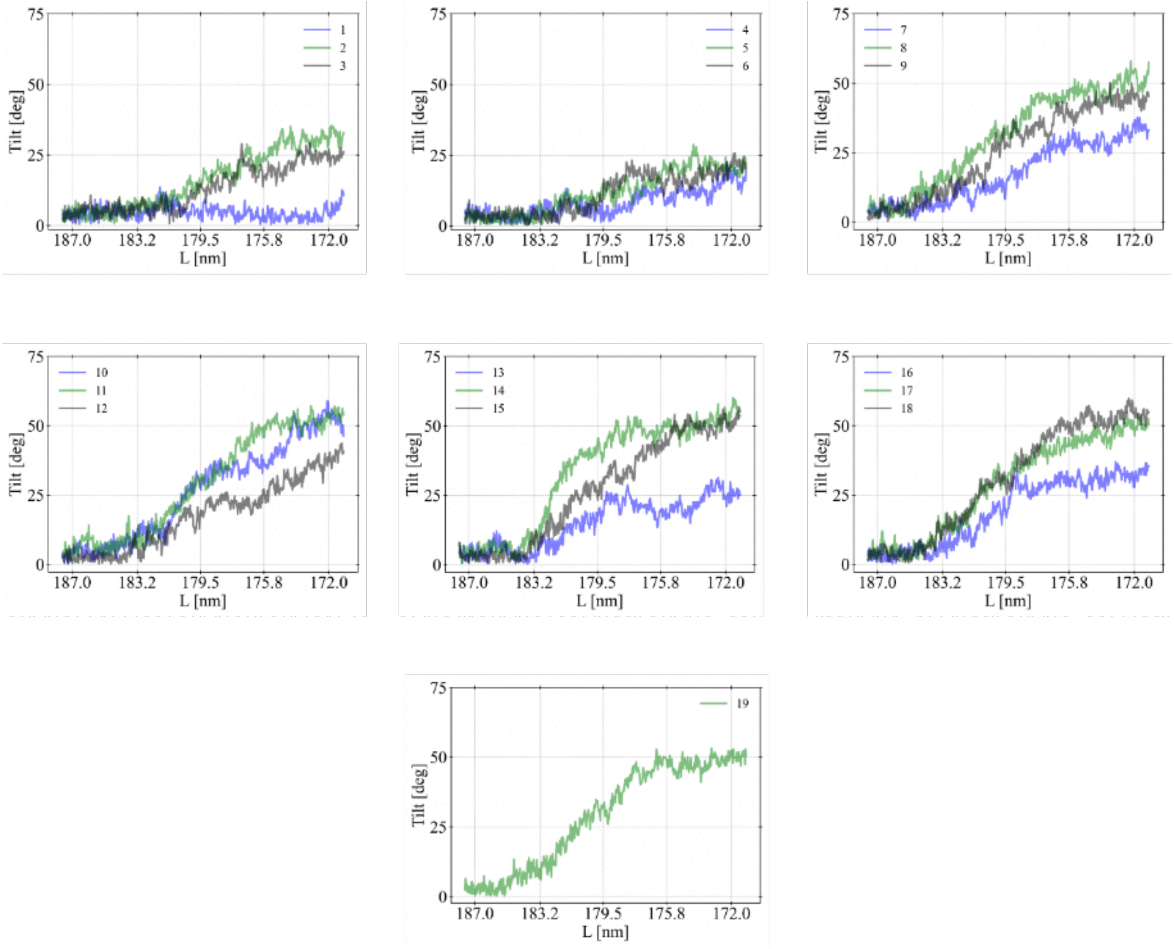


**Fig. S15. Orientational fluctuations of the 19 CAV1-8S complexes during compression.** Tilting of specific complexes increases substantially with the size of the invagination.

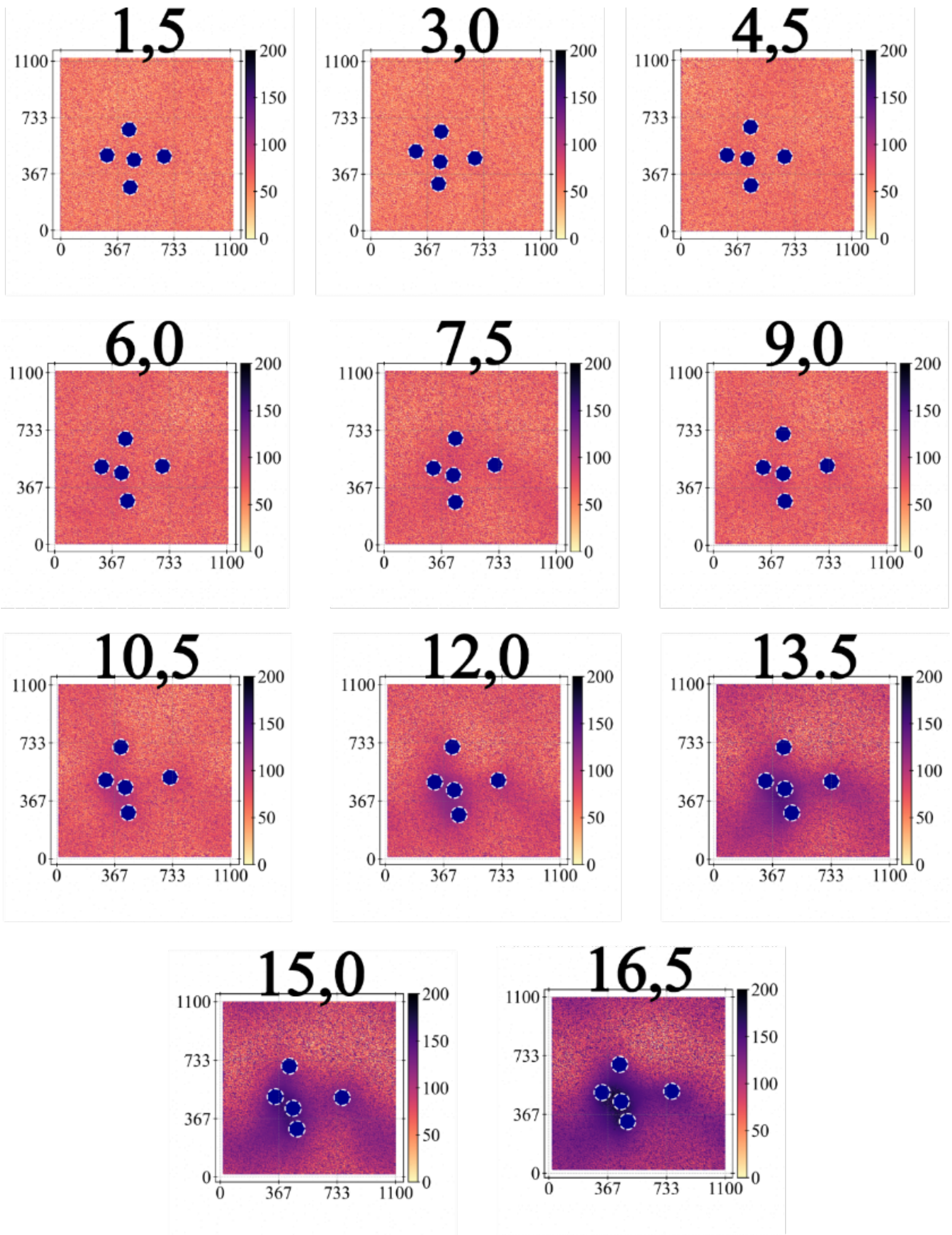


**Fig. S16. Height profiles for the system with 5 CAV1-8S complexes under compression computed with Gaussian Process Regression**. Numbers on top of the profiles specify the timestamp of the simulation in units of 1mio.

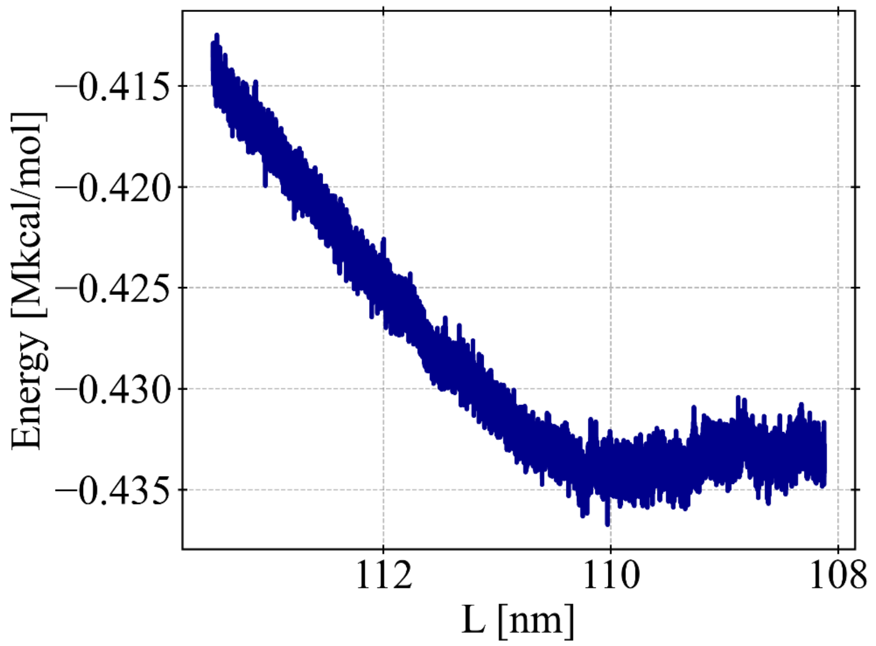


**Fig. S17. Energy landscape recorded during compression of the system with 5 CAV1-8S complexes.** Note that the profile is consistent with the simulation of 19 complexes.

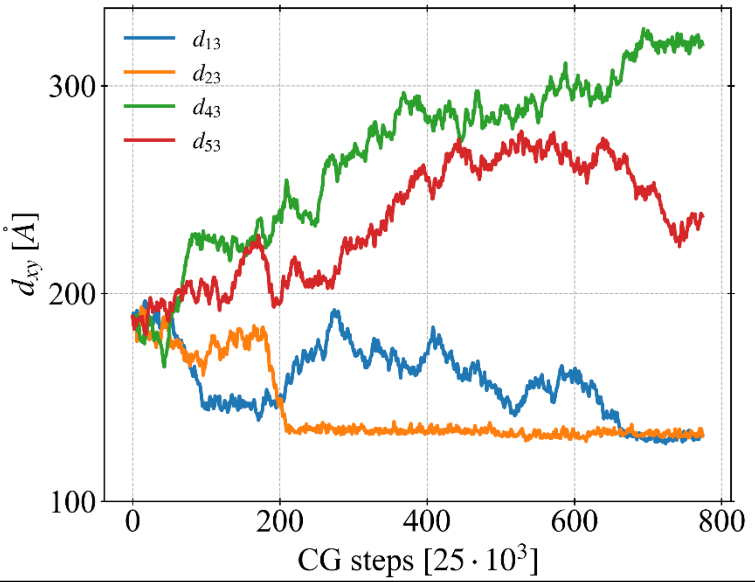


**Fig. S18. Inter-complex distances recorded from the simulation with 5 CAV1-8S complexes.** Distances of four complexes to the centrally positioned one are plotted. This shows that two complexes bind to and two other diffuse away from the central complex.

### 4. Movies

**Movie S1. Compilation of the REM-training of the CG-model.** Iterative optimization of the parameters of the potential energy converges to a correct binding configuration of the CAV1-8S complex in the membrane.

**Movie S2. Self-assembly of the lipid bilayer.** The developed CG-model accurately captures self-assembly of the lipid-bilayer (POPC head-groups are shown in green).

**Movie S3. Formation of a membrane invagination (side view).** This movie is a compilation of the simulation with 19 CAV1-8S complexes (under compression).

**Movie S4. Formation of a membrane invagination (top view).** This movie is a compilation of the simulation with 19 CAV1-8S complexes (under compression).
